# ALG-1, a microRNA argonaute, promotes vulva induction in *C. elegans*

**DOI:** 10.1101/2024.09.29.615693

**Authors:** Sunny Zhen, Christian E. Rocheleau

**Affiliations:** Department of Biomedical Sciences, University of Waterloo, Waterloo, ON, Canada; Division of Endocrinology and Metabolism, Department of Medicine, McGill University, Montreal, QC, Canada; Metabolic Disorders and Complications Program, Centre for Translational Biology, Research Institute of the McGill University Health Centre, Montreal, QC, Canada

## Abstract

Signaling by the LET-60 Ras GTPase/MPK-1 Extracellular Regulated Kinase pathway specifies the vulva cell fate in *C. elegans*. The *let-7* miRNA family negatively regulates LET-60 Ras but other miRNAs can also modulate vulva induction. To determine the impact of globally reducing miRNA function on LET-60 Ras-mediated vulva induction we analyzed the effect of loss of the ALG-1 miRNA regulator on vulva development. Contrary to our expectations, we find that ALG-1 promotes vulva induction independently of LET-60 Ras. We found that the reduced vulva cell fate induction of *alg-1* deletion mutants could be due to delayed development of the vulva, or a requirement to maintain the competence of the uninduced precursor cells.

## Description

LET-60 Ras is an essential component of an Epidermal Growth Factor Receptor/Ras GTPase/Mitogen Activated Protein Kinase that induces three of six epithelial cells to adopt vulva cell fates (Figure 1A) (Beitel *et al*., 1990; Han *et al*., 1990; Han and Sternberg, 1990; Schmid and Hajnal, 2015). During the second larval stage, six epithelial cells (P3.p-P8.p) are maintained in a competent state to be induced by a combination of LET-23 EGFR and LIN-12 Notch signaling pathways in the third larval stage (Greenwald *et al*., 1983; Yochem *et al*., 1988; Aroian *et al*., 1990; Wang and Sternberg, 1999). Three cells, P5.p, P6.p and P7.p, are induced by graded LIN-3 EGF-like signal from the overlying gonad (Figure 1A) (Sternberg and Horvitz, 1986; Hill and Sternberg, 1992; Katz *et al*., 1995). P6.p, the closest to the ligand is induced to adopt the primary (1°) vulval fate and lateral LIN-12 Notch signaling from P6.p paired with the graded LIN-3 signaling specifies the secondary (2°) vulval fates in P5.p and P7.p. P5.p, P6.p and P7.p undergo the stereotypical three rounds of division to produce the 22 nuclei that make up the mature vulva (Figure 1A, B top). The remaining P3.p, P4.p and P5.p cells adopt the tertiary (3°) non-vulval fate in which they divide once and fuse with the surrounding hypodermis, except for P3.p which 50% of the time fuses prior to dividing and adopts a fused (F) fate (Sternberg and Horvitz, 1986; Eisenmann *et al*., 1998).

**Figure 1.**
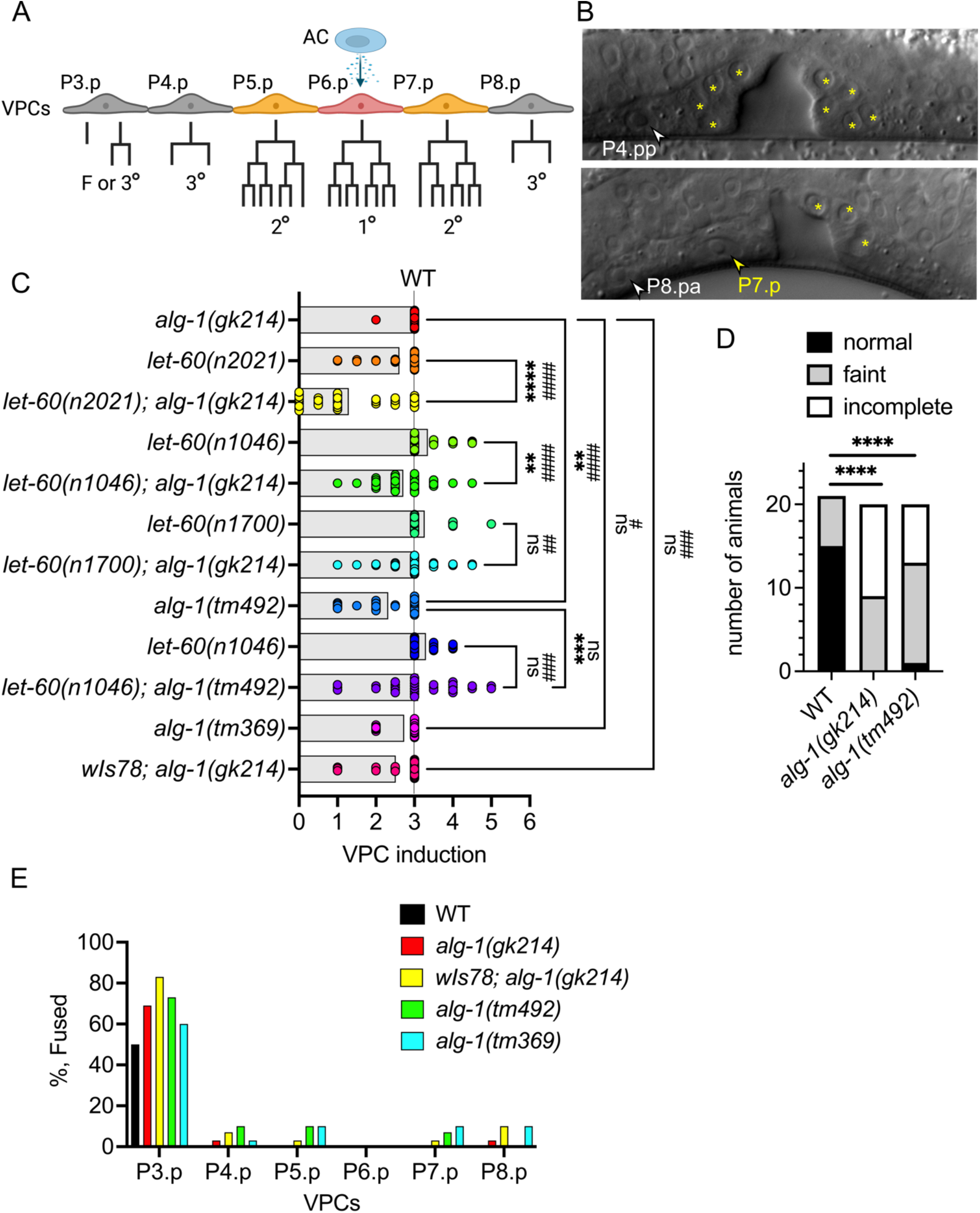
*alg-1* deletion alleles cause a partial vulvaless phenotype. (**A**) Schematic of vulva induction. Six vulva precursor cells (VPCs) P3.p-P8.p are competent to be induced to adopt vulval cell fates. However, only the VPCs closest to the anchor cell (AC) are induced to adopt 1° and 2° vulval fates which undergo three rounds of cell division to generate 22 vulval cells. The remaining VPCs adopt the 3° non-vulval fate and divide once with the exception of P3.p, which 50% of the time loses competence and adopts the fused (F) fate without dividing. Created using Biorender (**B**) Differential interference contrast images of normal (top) and partially induced (bottom) vulvas of L4 stage *alg-1(tm369)* deletion mutant strain. In the top image (anterior is left, ventral is down) yellow asterisks mark the visible nuclei of P5.p and P7.p descendants. White arrow marks the uninduced posterior daughter of P4.p. In the bottom image (anterior is right) yellow asterisks mark the visible nuclei of the descendants of P5.p and a yellow arrow marks the P7.p cell that failed to divide and appears to have adopted a fused fate. White arrow marks the uninduced anterior daughter of P8.p. (**C**) Bar graph/scatter dot plot depicting vulva induction in various *alg-1* and *let-60* mutants. Each dot represents the VPC induction score for one animal and the bar represents the average VPC induction score for the strain. A score of 3 is wild type (WT), <3 is vulvaless, and >3 is multivulva. Since the VPCs are still competent to be induced after the first division, half inductions are common when signaling is compromised. A one-way ANOVA with multiple comparisons was used to determine the significance between average VPC inductions scores \*\**P<0*.*01*, \*\*\**P<0*.*001* and \*\*\*\**P<0*.*0001*. A Fisher’s exact test was used to determine the significance of the vulvaless phenotypes ##*P<0*.*01*, ###*P<0*.*001* and ####*P<0*.*0001. ns*, not significant. (**D**) Bar graph depicting the number of animals with normal alae or defective alae that were either faint or incomplete in wild type, *alg-1(gk214)* and *alg-1(tm492)*. A Fisher’s exact test was used to determine the difference in normal versus defective (faint + incomplete) alae between wild type and the *alg-1* mutants. \*\*\*\**P<0*.*0001*. (**E**) Bar graph depicting the % Fused fate adopted by each VPC in the *alg-1* mutants compared to that expected for wild type. Between 30-34 animals were scored for the vulval phenotypes depicted in panels C and E.

Argonaute proteins are the core component of the miRISC complex that guide miRNAs to the 3’UTR of their mRNA targets to inhibit translation and/or initiate mRNA degradation (Frederick and Simard, 2022; Ambros, 2024). The *C. elegans* miRNA argonautes, ALG-1 and ALG-2, are redundantly required for embryogenesis (Grishok *et al*., 2001; Vasquez-Rifo *et al*., 2012) however, only *alg-1* deletions have post-embryonic miRNA associated phenotypes suggesting that ALG-1 has a greater role in regulating miRNA function during larval development.

The 3’UTR of *let-60 ras* has several *let-7* miRNA target sites (Johnson *et al*., 2005). Overexpression of *let-7* family members *mir-48* and *mir-84* partly suppress the multivulva phenotype of activating alleles of *let-60* (Johnson *et al*., 2005; Li *et al*., 2005). Furthermore, *mir-48* and *mir-84* have reciprocal expression patterns with *let-60* in the VPCs and this is dependent on the 3’UTR of *let-60* (Esquela-Kerscher *et al*., 2005; Johnson *et al*., 2005). Although, loss of *let-7* family members *mir-48, mir-84* and *mir-241* independently or in combination, fail to cause a vulval phenotype (Abbott *et al*., 2005), whereas precocious expression of *let-7* results in a multivulva phenotype (Hunter *et al*., 2013). Additionally, other components of the Ras/MAPK pathway have predicted *let-7* miRNA target sites (Johnson *et al*., 2005), and other miRNAs can differentially modulate the *let-60 ras* multivulva phenotype (Brenner *et al*., 2012). Thus, multiple miRNAs may regulate LET-60 Ras signaling at multiple targets.

To assess the global requirements of miRNAs in LET-60 Ras-mediated vulva induction, we tested for genetic interactions between a deletion allele of *alg-1, gk214*, with hypomorphic and hypermorphic alleles of *let-60 ras* (Ferguson and Horvitz, 1985; Beitel *et al*., 1990; Ding *et al*., 2005). While the *alg-1(gk214)* strain had close to normal vulva development, we found that it enhanced the vulvaless phenotype of *let-60(n2021)* hypomorphic allele, opposite what we would expect with a loss of the *let-7* miRNA family (Figure 1C). Next, we tested if *alg-1(gk214)* could suppress the multivulva phenotype of the *let-60(n1046)* hypermorphic mutant. Surprisingly, the *let-60(n1046); alg-1(gk214)* multivulva phenotype was not significantly suppressed, but instead displayed a vulvaless phenotype not seen in either single mutant (Figure 1C). Suspecting that one of the strains could have a genetic modifier in the background, we crossed *alg-1(gk214)* into the *let-60(n1700)* hypermorphic mutant that possess the same G13D point mutation found in the *let-60(n1046)* mutant (Beitel *et al*., 1990). Again, we saw a vulvaless phenotype in *let-60(n1700); alg-1(gk214)* double mutants (Figure 1C). We then crossed another deletion of *alg-1, tm492*, with *let-60(n1046)* (Yigit et al., 2006). Here we found that both *alg-1(tm492)* and *let-60(n1046); alg-1(tm492)* had similar vulvaless phenotypes suggesting that loss of *alg-1* itself results in a previously undescribed vulvaless phenotype (Figure 1C). Thus, we looked at a third deletion allele, *alg-1(tm369*) (Yigit et al., 2006), and an *alg-1(gk214)* strain which was further crossed with a seam cell marker, *wIs78* (Koh and Rothman, 2001). We found that both strains displayed a vulvaless phenotype that together, were significantly more vulvaless than the starting *alg-1(gk214)* strain (Figure 1B,C). Thus, the starting strain likely harbors one or more genetic suppressors that were lost in subsequent crosses with the *let-60* mutants and *wIs78*. These suppressors could be specific to VPC induction as we did not find differences in alae defects between the starting *alg-1(gk214)* strain and the *alg-1(tm492)* mutant (Figure 1D).

All VPCs divide once, whether they are induced or not induced, excluding P3.p which fuses with the surrounding hypodermis prior to induction about 50% of the time in wild-type hermaphrodites (Figure 1A). We found that in the *alg-1* deletion alleles, a low level of VPCs failed to divide suggesting that either they fused prematurely and lost competence or had a cell cycle defect (Figure 1E). While all *alg-1* mutants had over 50% of P3.p cells adopting the F fate, only *wIs78; alg-1(gk214)* reached a statistically significant increase above 50%. However, the low level of fused VPCs does not account for the vulvaless phenotype seen in *alg-1* mutants as most uninduced cells divided once. Another phenotype we noted was a low percentage of incompletion lineages in *let-60(gf); alg-1(-)* strains, in which the induced VPCs failed to complete the third cell division by the fourth larval stage, suggesting either a delayed or incomplete execution of the lineages.

Here, we report that *alg-1* mutants have a vulvaless phenotype. Mutations in *alg-1* and *ain-1 GW182*, a component of the miRISC complex, were previously identified as suppressors of the *lin-31* multivulva phenotype (Ding *et al*., 2005; Morita and Han, 2006); however, the role of *alg-1* in regulating vulval cell fate induction was not characterized. While the *alg-1* mutant vulvaless phenotype is inconsistent with a simple loss of *let-7* family miRNAs antagonizing *let-60 ras*, the incomplete lineages could reflect a heterochronic phenotype (Euling and Ambros, 1996). Mutations in the *lin-4* miRNA cause a vulvaless phenotype with a failure of division in some VPCs and supernumerary divisions in other VPCs (Chalfie *et al*., 1981; Sulston and Horvitz, 1981; Ambros and Horvitz, 1984; Ferguson and Horvitz, 1985). The *mir-48 mir-241; mir-84* triple mutant in combination with the *lin-46* heterochronic mutant has a retarded vulva development phenotype (Abbott *et al*., 2005), and *alg-1(gk214)* enhances the retarded vulva development phenotype of a *lin-66* heterochronic mutant (Morita and Han, 2006). Thus, some of the *alg-1* vulval phenotypes could be a result of heterochronic phenotypes due to a reduction in *lin-4* and *let-7* miRNA family function. However, the *lin-4* and *mir-61* miRNAs promote LIN-12 Notch signaling and the 2° vulval fate (Yoo and Greenwald, 2005; Li and Greenwald, 2010). Notably, we found that P6.p was always induced in the *alg-1* deletions, which could be consistent with a specific failure to induce the 2° vulval fates or could reflect a partial loss of LET-60 Ras signaling as P6.p was induced in all hypomorphic *let-60(n2021)* mutant animals analyzed. The failure of P6.p to adopt an F fate is also reminiscent of some Wnt signaling mutants (Myers and Greenwald, 2007). Thus, ALG-1 and various miRNAs could promote vulva cell fate induction via regulation of developmental timing and promoting the LIN-12 Notch, LET-60 Ras, and Wnt signaling pathways. Transcriptomic analysis of the VPCs identified transcripts predicted to be regulated by 114 miRNAs (Zhang *et al*., 2022). Further work will be required to delineate the mechanisms and miRNAs promoting vulva induction with ALG-1. The partial vulvaless phenotype of *alg-1* mutants could be leveraged for the further identification and characterization of miRISC regulators.

## Methods

Strain construction and maintenance were done as previous described (Stiernagle, 2006; Fay, 2013). N2 was the wild type strain from which all mutants were derived. HB101 *E. coli* was used as a food source. Wormbase.org was critical for identifying reagents and designing experiments (Fay, 2013). Vulva induction scoring was done as previously described (Gauthier and Rocheleau, 2017). Differential interference contrast images were acquired on an Axio Imager A1 using the Axiocam 305 mono camera with Zen software (Zeiss, Germany). Graphpad Prism 10 was used to generate graphs and statistical analyses.

### Reagents

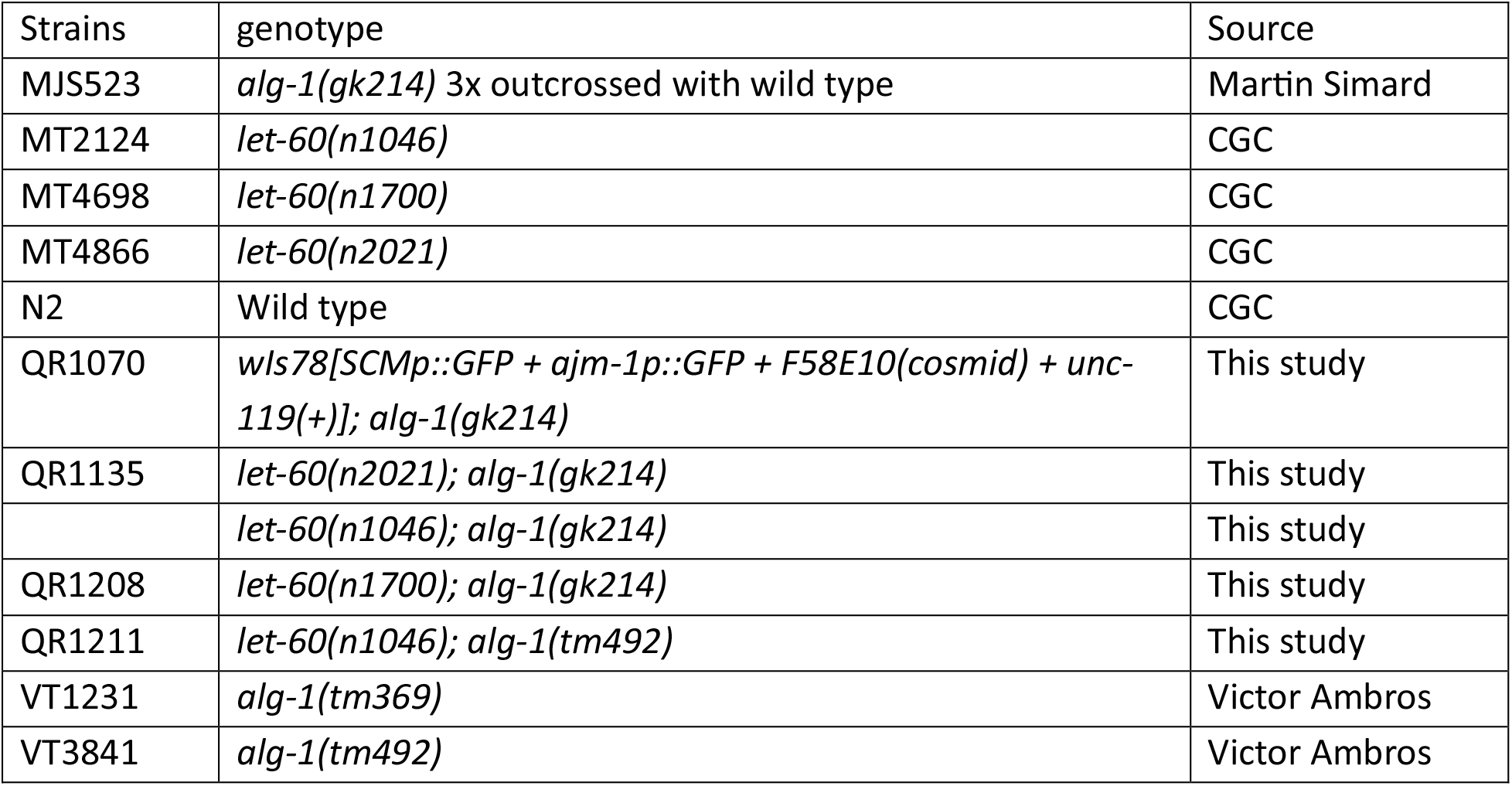

## Acknowledgements

We thank Jung Hwa Seo for technical assistance. We thank the Simard (Université Laval) and Ambros (University of Massachusetts, Worcester) for sharing strains. Deletion alleles of *alg-1* were generated by the National Bioresource Project for the nematode (Tokyo Women’s Medical University School of Medicine, Japan) and the International *C. elegans* Gene Knockout Consortium (Oklahoma Medical Research Foundation, USA and the University of British Columbia, Canada). Some strains were provided by the Caenorhabditis Genetics Center (CGC), which is funded by NIH Office of Research Infrastructure Programs (P40 OD010440).

## Contributions

